# Efficient Correction of the Sickle Mutation in Human Hematopoietic Stem Cells Using a Cas9 Ribonucleoprotein Complex

**DOI:** 10.1101/036236

**Authors:** Mark A. DeWitt, Wendy Magis, Nicolas L. Bray, Tianjiao Wang, Jennifer R. Berman, Fabrizia Urbinati, Denise P. Muñoz, Donald B. Kohn, Mark C. Walters, Dana Carroll, David K. Martin, Jacob E. Corn

**Author notes:** Co-corresponding authors: Jacob Corn, David Martin and Dana Carroll.

## Abstract

Sickle Cell Disease (SCD) is a serious recessive genetic disorder caused by a single nucleotide polymorphism (SNP) in the ß-globin gene (***HBB***). Sickle hemoglobin polymerizes within red blood cells (RBCs), causing them to adopt an elongated “sickle” shape. Sickle RBCs damage vasculature, leading to severe symptoms, ultimately diminishing patient quality of life and reducing lifespan. Here, we use codelivery of a pre-formed Cas9 ribonucleoprotein complex (RNP) and a singlestranded DNA (ssDNA) oligonucleotide donor to drive sequence replacement at the SCD SNP in human CD34+ hematopoietic stem/progenitor cells (HSPCs). Corrected HSPCs from SCD patients produce less sickle hemoglobin protein and correspondingly increased wild-type hemoglobin when differentiated into erythroblasts. When injected into immunocompromised mice, treated HSPCs maintain editing long-term at therapeutically relevant levels. These results demonstrate that the Cas9 RNP/ssDNA donor approach can mediate efficient HSPC gene editing and could form the basis for treatment of SCD by autologous hematopoietic cell transplantation.

## Introduction

Sickle Cell Disease (SCD) is a recessive genetic disorder that afflicts at least 90,000 predominantly African-American individuals in the US and hundreds of thousands worldwide^1,2^. The genetics and molecular basis of SCD have been understood for almost seventy years, but interventions that address the underlying cause of the disease have lagged behind^3,4^. SCD is caused by a single nucleotide polymorphism (SNP) in the seventh codon of ß-globin (***HBB***), one of two globins comprising the major adult form of hemoglobin. The resulting sickle glutamate-to-valine substitution renders hemoglobin prone to polymerization, leading to characteristic “sickle” shaped red blood cells (RBCs). Sickle RBCs have a markedly reduced persistence in the bloodstream and damage vasculature. Major clinical manifestations of SCD are chronic anemia, severe pain episodes, and progressive damage to multiple organs. The disease causes a 30-year decrement in lifespan, and a greatly diminished quality of life^2,5-7^.

A broadly applicable cure for SCD remains elusive. RBCs are produced from hematopoietic stem cells (HSCs) in the bone marrow, and autologous hematopoietic cell transplantation (HCT) from an unaffected HLA-matched donor is currently the only cure for SCD^8,9^. However, HCT has been employed sparingly because of the difficulty in identifying donors within the affected populations, along with risks associated with the intensive transplant procedure (requiring ablative chemotherapy and immune suppression), including potentially fatal graft-versus-host disease^9,10^.

Gene editing has recently emerged as a promising avenue to treat hematopoietic genetic disease^11-13^. ***Ex vivo*** editing of autologous hematopoietic stem cells (HSCs) would be followed by reimplantation of edited cells, bypassing donor requirements and eliminating the need for immunosuppression. Because SCD is monoallelic and sickle RBCs have a markedly shorter circulating lifespan than wild type cells, relatively low levels of correction may still yield significant clinical benefit^14^. Indeed, observations after allogeneic HCT suggest that a significant benefit occurs when as few as 2% of HSCs carry normal HBB^15-17^. Still, correction of the SCD SNP at this level in HSCs has to date proved elusive^11,18^.

During gene editing, a targeted nuclease creates a double strand break (DSB) that can be repaired by one of two mechanisms: error-prone non-homologous end joining (NHEJ) that leads to genomic insertions and deletions (indels), or templated homology-directed repair (HDR) to precisely insert, delete, or replace genomic sequence^19^. The recent development of Cas9, a programmable RNA-targeted DNA endonuclease, has ignited an explosion of interest in gene editing to potentially cure many genetic disorders, including SCD^20,21^. Guided by a single guide RNA (sgRNA), the Cas9 nuclease can be readily programmed to cut a target locus within the genome, allowing rapid iteration and optimization not possible with other gene editing approaches^20,22^.

Optimized methods for efficient ***ex vivo*** gene editing of human HSCs are required to enable a CRISPR/Cas9-based treatment for blood disorders such as SCD. Recent work has demonstrated that Cas9 can be used for ***in vitro*** reversion of the SCD mutation in laboratory cell lines^23,24^ and induced pluripotent stem cells (iPSCs)^25^, as well as efficient knockout of an erythroid enhancer in an immortalized cell line^26^ and ***HBB*** in HSCs^24^. Most recently, zinc finger nucleases (ZFNs) have been used to correct the SCD mutation in HSCs, albeit at levels less than 1% in the longterm re-populating stem cell population^18^. To date gene editing has yet to achieve long-term correction of the SCD mutation in re-populating HSCs at clinically relevant levels, as defined by the ability of corrected cells to persist in immunocompromised mice^11,18^.

Delivery of purified Cas9 protein and ***in vitro*** transcribed sgRNA as a preassembled ribonucleoprotein complex (RNP) via electroporation can be used for gene knockout in a variety of cell types, including primary cells, and we have recently extended this approach to enable extremely efficient HDR sequence replacement in laboratory cell lines^27-30^. Given the efficiency and speed of Cas9 RNP-based editing, we reasoned that the Cas9 RNP could form the basis for a protocol to achieve lasting correction of the SCD SNP in human HSCs.

We first used an erythroleukemia-derived cell line to exhaustively explore a panel of Cas9 RNPs that cut near the SCD SNP. We then used the most effective RNPs to develop editing methods in human HSPCs, achieving up to 32% HDR in HSPCs ***in vitro.*** We demonstrate efficient correction of the SCD mutation in SCD HSPCs, with corresponding production of WT adult hemoglobin (HbA) protein in edited erythroblasts. After engrafting edited HSPCs in immune-compromised mice, HDR-based gene editing at the SCD locus is maintained at clinically beneficial levels of editing several months post-engraftment.

## Results

### Prioritizing SCD editing reagents in a model cell line

Pairing Cas9 with different sgRNAs can lead to different activities towards the same gene target^31^. For HDR-mediated editing, each sgRNA must in turn be paired with an HDR donor template that encodes the desired nucleotide changes. We used short single-stranded DNA (ssDNA) donors that can be rapidly optimized, and co-delivered with the Cas9 RNP by electroporation. The rules governing ssDNA donor design are still unclear and may differ among cell types, though our laboratory has developed guidelines based on the mechanism of Cas9’s cleavage activity^30^. The electroporation protocols to best deliver a Cas9 RNP and associated donor template may also differ between cell types. Hence, discovering a suitable approach to edit the SCD allele in primary HSPCs represents a large, combinatorial problem, especially since HSPCs are costly to obtain and difficult to culture.

We searched for maximally active sgRNAs in K562 cells, a human erythroleukemia line that resembles early committed hematopoietic progenitors. Two key restrictions when designing sgRNAs and donors for HDR experiments are the distance between the sgRNA recognition site and the mutation, and the ability to silently ablate the sgRNA protospacer adjacent motif (PAM). This latter constraint ensures that Cas9 cannot re-cut corrected alleles, preventing the simultaneous introduction of indels and the template SNP.

To find maximally active sgRNAs, we transcribed ***in vitro*** a panel of all sgRNAs targeted within 100 bp of the SCD SNP for which PAMs could be silently mutated (termed **G1-G18**, Fig. **1A**)^23,24^. Some of these guide RNAs targeting HBB have previously been reported, but to the our knowledge have not been tested in a Cas9 RNP format^23^. We also truncated three of these sgRNAs at the 5’ end (yielding an 18-19 nt guide sequence), which has been reported to reduce off-target effects without compromising on-target efficacy^32^. We designed an initial ssDNA donor **(T1)** that contained silent mutations in the PAMs of all sgRNAs and a WT-to-SCD edit (K562 cells are WT at ***HBB*** Fig. **S1A**). The combination of the WT-to-SCD SNP mutation and removal of the **G5** PAM generates a silent SfcI site (SCD), allowing easy tracking of HDR-mediated editing (Fig. **S1A**).

**Figure 1.**
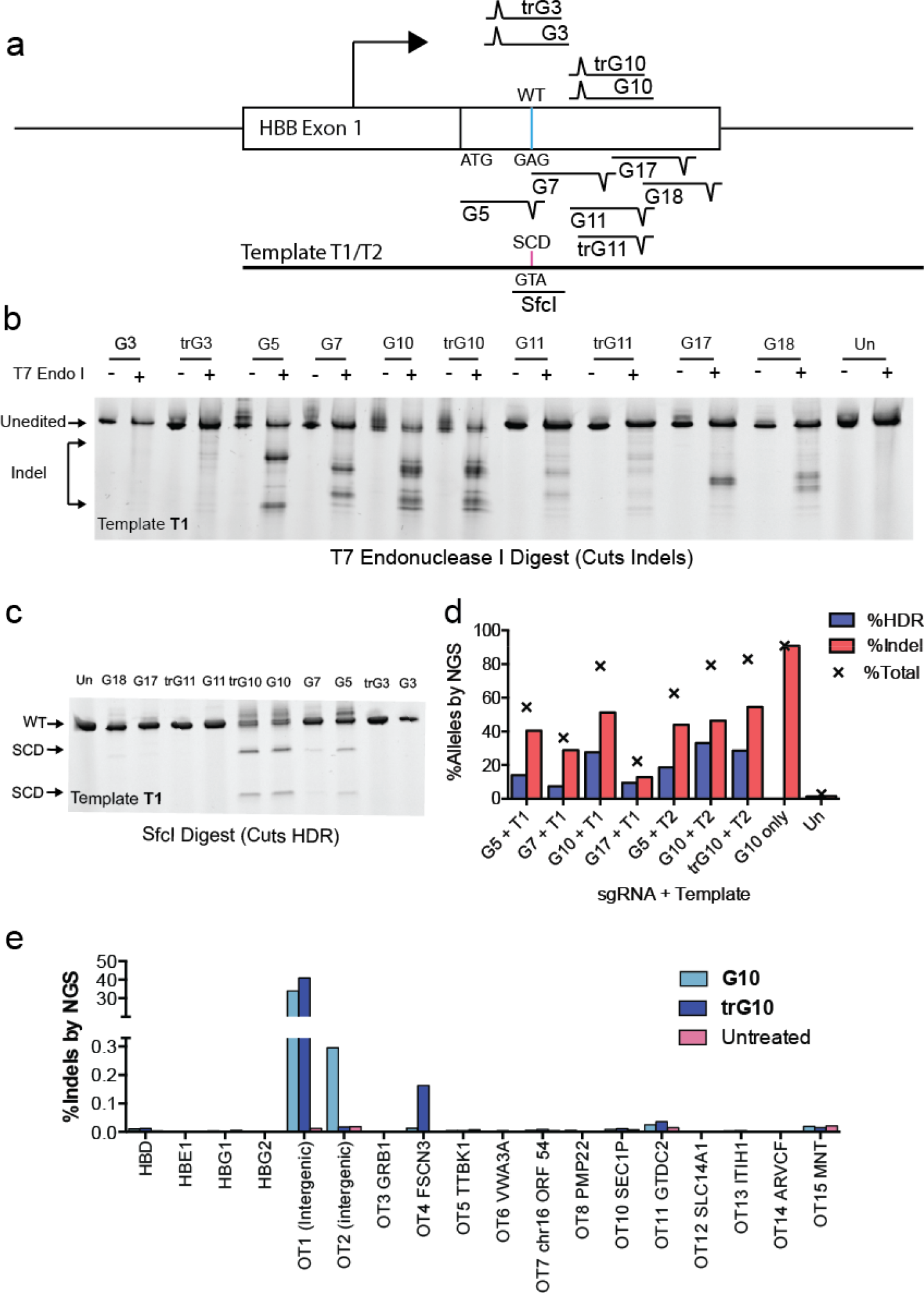
Editing the SCD SNP in K562 cells. A) Schematic depicting experimental approach to editing in K562 cells. A panel of 10 sgRNAs that cut within 50 bp of the SCD SNP was selected. A WT-to-SCD edit was programmed by an ssDNA template (Tl) bearing silent PAM mutations for the sgRNAs, which also introduces a Sfcl restriction site. B) T7 endonuclease 1 assay depicting indel formation in pools of cells edited by candidate RNPs. C) Editing of candidate sgRNAs detected by ***Sfcl*** digestion. **G5, G10** and a truncated variant, **trG10**, efficiently edit in K562 cells. D) Gene modification of select sgRNAs and templates at the SCD SNP, assessed by NGS. See Fig. S1 for definitions of donors **T1** and **T2.** E) Analysis of off-target cutting by the **G10** RNP at sites predicted by the online CRISPR-Design tool, in K562 cells, determined by NGS.

We assembled Cas9 RNPs from each candidate sgRNA and individually delivered them to K562 cells together with the ssDNA donor by electroporation. We used a “standard” dose of 100 pmol each RNP and template per 150,000 K562 cells, based on previous work^27^. A T7 endonuclease I digest of PCR amplicons from edited cell pools, which in this case detects mismatches arising from both NHEJ- and HDR-mediated repair of Cas9 DSBs, revealed that RNPs bearing only two sgRNAs **(G3** and **trG3)** were inactive at ***HBB*** (Fig. **1B**). Sfcl digest showed that three of these sgRNAs, **G5, G10**, and **trG10** (an 18 nt truncated form of **G10)** could produce HDR-mediated editing of the SCD SNP **(Fig. 1C).**

We quantified the frequency of HDR and indel formation at the SCD SNP using next-generation sequencing (NGS) of PCR amplicons derived from genomic DNA extracted from pools of edited cells (Fig. **1D**), and confirmed HDR rates in biological replicate by droplet digital PCR (ddPCR, Fig. **S2**). We used both the **T1** template and a derivative of **T1** bearing only the WT-to-SCD edit and the **G5** and **G10** PAM mutation **(T2).** Most candidate sgRNAs induced substantial quantities of indels (>40% of reads), and **G5, G10**, and **trG10** also yielded high levels of WT-to-SCD HDR (19% of reads for **G5**, 32% of reads for **G10)** from multiple donor designs. Using a phosphorothioate-protected ssDNA donor did not improve HDR frequencies (Fig. **S1B**). When the **G10** Cas9 RNP was provided without an ssDNA donor, 90% of reads contained indels, indicating excellent delivery of the RNP to K562 cells. The **G5** guide targets a sequence very similar to one found in the closely-related hemoglobin-delta (***HBD***) gene, and we experimentally verified that this guide induces indels in ***HBD*** (Fig. **S3**). Hence, we selected the truncated **trG10** guide for further testing.

To optimize an ssDNA donor for HDR, we drew guidance from recent work in our laboratory showing that upon binding its target, Cas9 releases the PAM-distal non-target strand, and that asymmetric homology arms taking advantage of this property can increase HDR efficiency^30^. For sgRNAs that target the sense strand such as **G10**, the best template to employ matches the sense strand (annealing to the antisense strand) and has a long 5’ homology arm and a shorter 3’ annealing arm (Fig **S4A**). Based on these principles, we will subsequently identify templates by the lengths of their 5’ and 3’ homology arms relative to the **G10** cut site. For example, the initial template **T2** will be called **T88-107.** Unless noted, all these templates bear only the **G5** and **G10** PAM mutations. We tested the effects of asymmetric templates containing the SCD-to-WT edit and the **G5** and **G10** PAM mutations (similar to **T2**, Fig. **S1**) on HDR-mediated editing in K562 cells (Fig. **S4B**). We found that a template with a 111 nt 5’ arm and a 57 bp 3’ arm **(Tlll-57)** yielded an HDR frequency of 35%, a modest improvement in efficiency over the original template, **T88-107.** High rates of HDR in all conditions, and a modest increase in HDR using asymmetric templates, were also confirmed by ddPCR (Fig. **S2**).

The ability of Cas9 to cut at off-target sites is a concern for the development of Cas9-based therapies^33,34^. While this tendency could be reduced by the use of Cas9 RNP delivery and/or truncated sgRNAs^27,32,33^, we used NGS of PCR amplicons to analyze the off-target activity of candidate Cas9 RNPs in K562 cells (Fig. **1E**). We selected candidate off-targets by sequence similarity using a popular off-target prediction tool^34^. We assessed the two top-scoring off-targets (which are both intergenic), the top 13 exonic off-targets, and the predicted off-target sites of the four globin genes (***HBD, HBE1, HBG1,*** and ***HBG2***, Fig. S5)^35^.

We observed very little exonic off-target activity, with most off-target sites showing no Cas9-dependent indel formation at rates within the limit of detection (∼0.001%). Substantial off-target activity was observed for both **G10** and **trG10** at a top-scoring intergenic site (0T1, chr9:104,595,865, 55% indel formation), which lies ∼3 kb from the nearest annotated genic sequence. High off-target activity by **G10** at this site has been observed previously^23^. Off-target activity was detected with **G10** at the other intergenic site (OT2, chr17:66,624,238, 0.30%), but this was absent with the truncated guide **trG10**. Low but detectable off-target activity was observed with both the **G10** and **trG10** RNPs at two exonic off-targets, FSCN3 and GTDC2. Indel formation was lower for the full-length **G10** RNP. (FSCN3: 0.013% and 0.16% indel for **G10** and **trG10** respectively; GTDC2: 0.025% and 0.035% indel for **G10** and **trG10**, respectively).

### Optimizing candidate Cas9 RNPs for efficient in vitro CD34+ HSC editing

While functional sgRNAs tend to be effective in multiple cell types, editing outcomes (NHEJ vs. HDR) differ across cell types. Indeed, protocols to ablate genes via knockout have been described for HSPCs and HSCs^13,36^, but HSCs have remained refractory to efficient HDR-based editing^11,18^. We used the trG10 sgRNA, which mediates efficient HDR and has few off-targets in K562 cells, to re-optimize editing in mobilized peripheral blood CD34+ HSPCs. Since these cells were obtained from healthy wild-type (WT) donors, our initial experiments used ssDNA donor templates bearing a WT-to-SCD mutation.

We first re-optimized electroporation conditions, template design, and RNP/donor template dose in HSPCs. To selectively interrogate editing in viable cells, we assayed edited HSPCs cultured for 7 days after editing in erythroid expansion conditions (erythroid-expanded), along with edited HSPCs cultured for only 2 days (un-expanded). Editing outcomes were analyzed qualitatively by T7E1 digest (indels and HDR), Sfc1 restriction analysis (HDR), and quantitatively by NGS. In HSPCs, we found that an asymmetric HDR template that was most effective in K562 cells **(T111-57)** drove HDR more efficiently at a lower Cas9 dose of 100 pmol RNP per 150,000 HSPCs, while a shorter template (111 nt left, 27 nt right) was more efficient at a higher dose of Cas9 (200 pmol RNP per 150,000 HSCs) (Fig. **S6**-**S7**). Finally, we tested the effects on editing in HSPCs of Scr7, an NHEJ inhibitor shown to increase HDR-mediated repair in other cell types, on editing in HSPCs, but did not observe a consistent increase in HDR (Fig. **S6**).

Guided by this re-optimization, we used NGS to analyze editing in HSPCs at the SCD SNP using multiple templates, doses of RNP, and electroporation conditions (Fig. **2A**). We achieved efficient total editing in HSPCs (%indel + %HDR = 50-80% of alleles), indicating good delivery of the **trG10** RNP to HSPCs. In erythroid-expanded HSPCs, we observed up to 21% HDR at the SCD SNP, with rates generally higher than 10%. We observed higher HDR frequencies in un-expanded HSPCs, with up to 21% of alleles converted at low dose of Cas9, and 32% alleles converted at high dose (Fig. **2A**). In general, higher editing was accompanied by apparent reduction in viability as measured by fewer viable cells remaining post-treatment (data not shown), consistent with neutral or deleterious selection within the edited cell population. Indels typically represented ∼40% of alleles. In separate experiments, high rates of HDR were confirmed in biological replicate by ddPCR (Fig. **S8**). These experiments establish efficient ***in vitro*** editing of CD34+ HSPCs using the Cas9 RNP, including scarless HDR-mediated sequence replacement.

**Figure 2.**
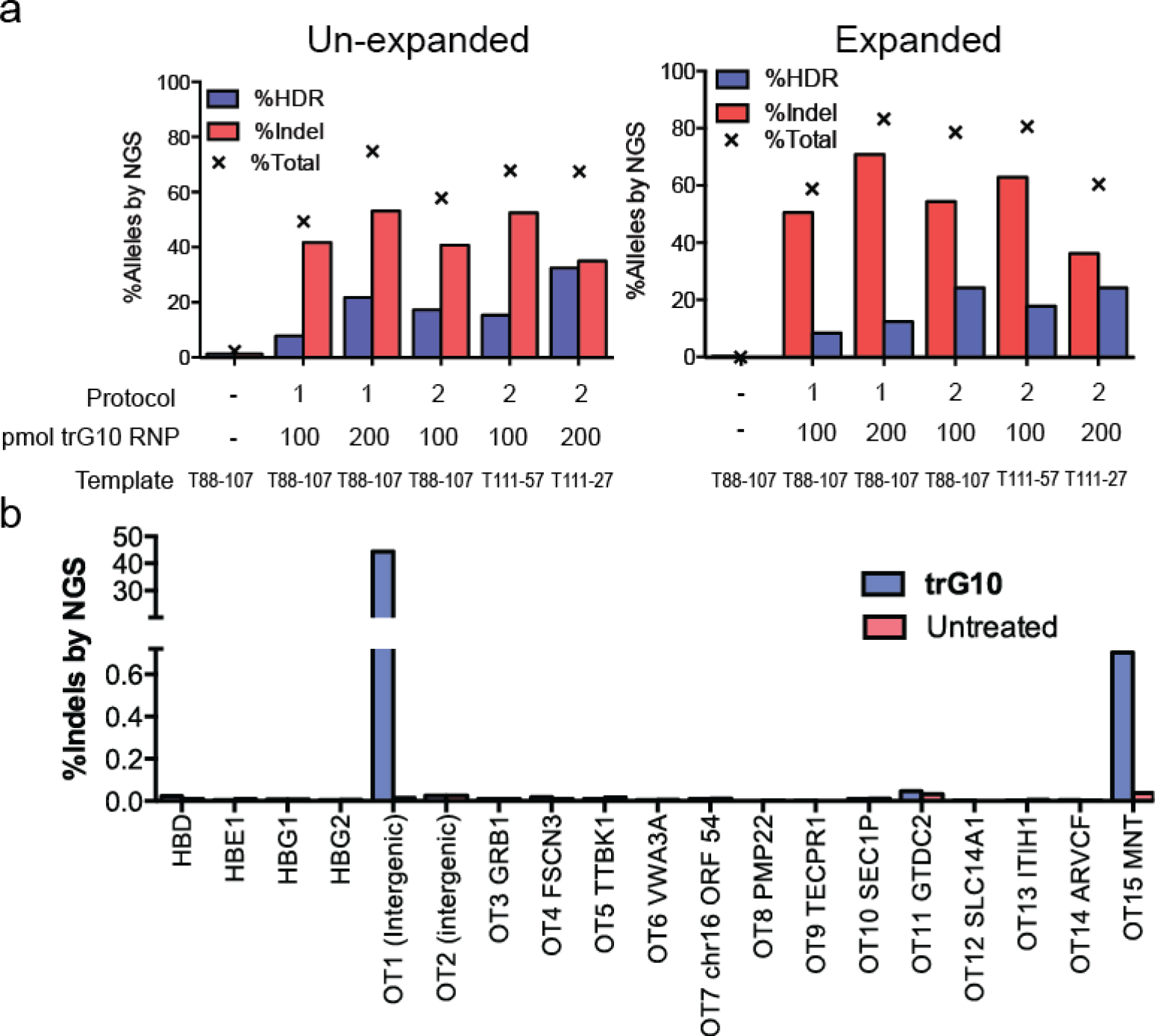
Editing of wild-type human CD34+ HSPCs by the Cas9 RNP. A) Analysis of editing in un-expanded HSPCs (left) and erythroid-expanded HSPCs (right), using **trG10** RNP and conditions as indicated. Templates, which are asymmetric about the **G10** cut site, are designed as described in the text. B) Indel formation at off-target sites in HSPCs edited with the **trG10** RNP, compared to untreated cells. Targets were selected using the online CRISPR-design tool.

We analyzed off-target activity by the **trG10** RNP in HSPCs using identical target selection criteria as for K562 cells (Fig. **2B**). Most predicted off-targets showed no detectable indel formation at genic off-targets, although cutting at the previously observed intergenic site remained high (0T1, 44% indel). When offtarget cleavage was observed in HSPCs, it frequently did not correspond with observations in K562 cells. For example, some indels were observed at FSCN3 and GTDC2 as before, but at a reduced rate (<0.05% in HSPCs vs. up to 0.16% in K562). Off-target activity in HSPCs was also observed at the MNT transcriptional repressor (0.7% indel), which was not seen in K562 cells, but the indels are located in the 3’ untranslated region, approximately 2 kb downstream from the stop codon. The biological ramifications of off-target activity at the MNT 3’ UTR warrant further analysis prior to clinical application of the **trG10** RNP.

### Efficient correction of the SCD mutation in HSPCs leads to production of wild type and fetal hemoglobin protein

Our success in WT-to-SCD editing HSPCs implies that the same method could be used to edit SCD HSPCs to WT. We obtained CD34+ HSPCs from de-identified SCD patient whole blood discard material after an exchange transfusion. We corrected the mutation using the **trG10** RNP and ssDNA donors programming an SCD-to-WT edit. These SCD-to-WT templates, which are denoted by the suffix **"S”**, encode the same number of mutations as the WT-to-SCD templates, with the base identity different only at the SCD SNP. Measuring editing by both NGS and ddPCR, we found that the SCD HSPCs were corrected at levels similar to those observed in WT HSPCs from mobilized blood, with up to 25% of alleles edited to WT at high RNP dose and 18% corrected at low RNP dose (Figs. **3A**, **S9).**

**Figure 3.**
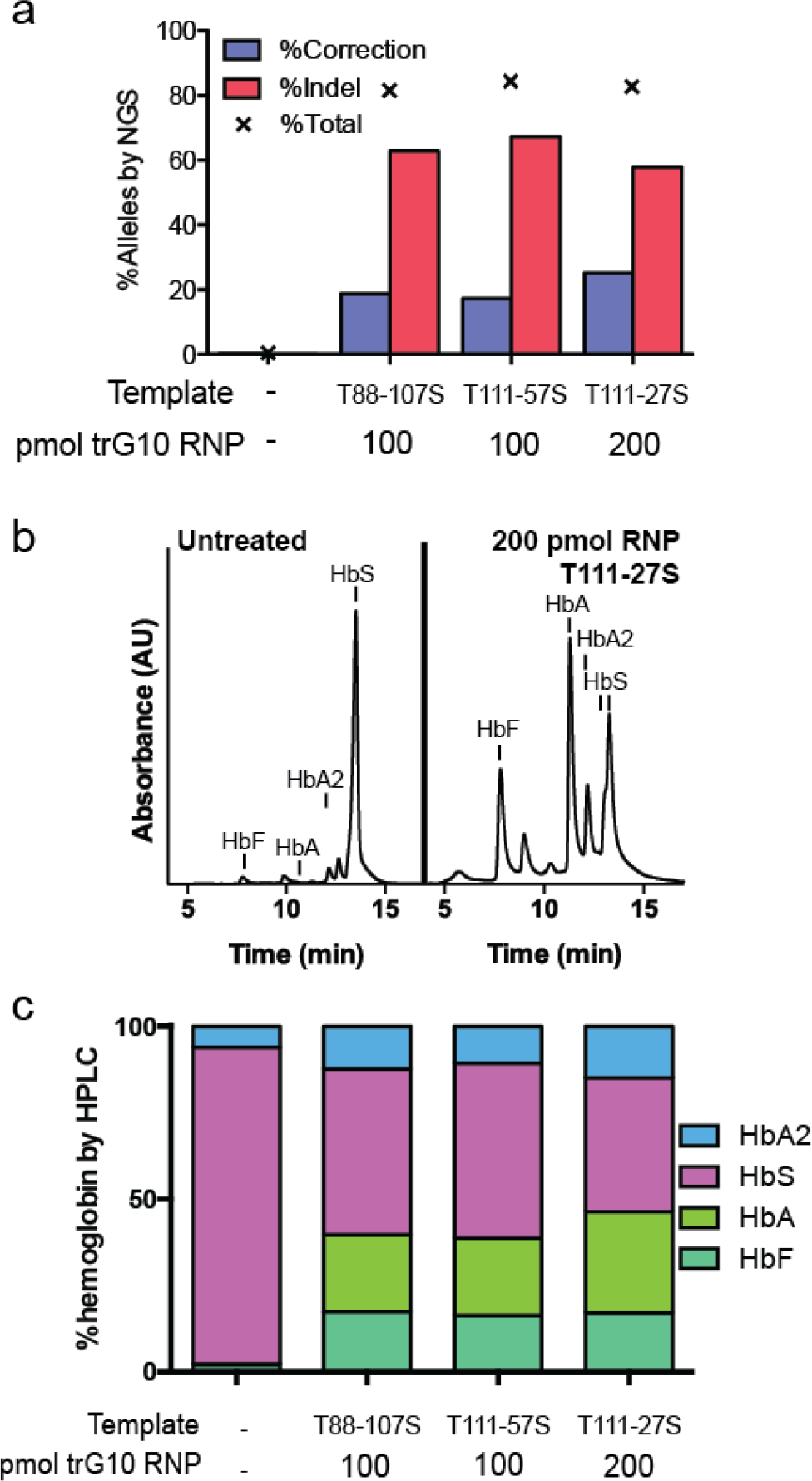
Correction of the SCD mutation in SCD HSPCs. A) Editing the SCD mutation in un-expanded CD34+ HSPCs from the whole blood of SCD patients, assessed by NGS. B) HPLC trace depicting hemoglobin production in SCD HSPCs edited with 200 pmol of the trGiO RNP and the T111-27S donor, compared to untreated HSPCs, after differentiation into erythroblasts. Significant increases in HbA, HbF, and HbA2 are apparent. C) Stacked bars showing HPLC results, with HSPCs edited as indicated prior to differentiation into erythroblasts.

To analyze the hemoglobin production potential of corrected HSPCs, we differentiated pools of SCD-to-WT edited HSPCs into enucleated erythrocytes and late-stage erythroblasts and measured the production of hemoglobin variant proteins by HPLC^18,37,38^. We found that corrected HSPCs produce substantial amounts of HbA, with a concomitant decrease in sickle hemoglobin (HbS) (22.2%-22.4% HbA at low dose RNP, 29.3% HbA at high dose RNP, Fig. **3B**-**3C**). Interestingly, we observed a substantial increase in fetal hemoglobin (HbF) in edited cells, far greater than the relative decrease in HbS (16.3%-17.4% HbF in edited cells vs 2.0% HbF in unedited cells, but 38.7% HbS in edited cells compared to 92.1% HbS in unedited cells). This may reflect a relative increase in γ-globin (fetal ß-like globin) production in cells that have lost production of ß-globin through induction of indels. Furthermore, the indels generated by the **trG10** RNP contain a large fraction (20-30%) of 9 bp in-frame deletions (Fig **S10**), suggesting the possible production of an altered sickle-like hemoglobin.

### Edited HSPCs repopulate in vivo and maintain therapeutically relevant levels of editing

The **trG10** Cas9 RNP paired with an appropriately designed HDR donor template can mediate efficient ***in vitro*** editing of the SCD SNP in CD34+ HSPCs. But ***in vitro*** experiments do not interrogate the efficiency of editing in long-term repopulating stem cells, which represent a small minority of CD34+ cells^39^. While it is possible to enrich more primitive stem cells using markers such as CD9 0 ^39,40^, a definitive means of assaying re-populating stem cells is through long-term xenografting in an immunodeficient mouse model, such as the NOD/SCID/IL-2rγ^nul1^ (NSG) mouse^41^. In this model, human cells must engraft in the mouse and edited cells must persist over several months at therapeutically relevant levels (1-5% correction), after which time all the human cells in the bone marrow are derived from re-populating true HSCs that mirror the proportion of editing at engraftment^18^’^41^.

To assess editing of the SCD SNP in re-populating stem cells, we injected pools of WT CD34+ HSPCs edited with the **trG10** RNP and the **T88-107** template into NSG mice^41^. Because HDR is down-regulated in G1 and G0 and up-regulated in the S and G2 phases of the cell cycle^42,43^, we also tested whether HDR rates could be improved by adding cytokines to the culture medium to stimulate proliferation of hematopoietic progenitors three days prior to editing, thereby promoting entry into the cell cycle (“stimulated HSPCs”). We then compared editing in these cells pre- and post-engraftment to cells maintained for only one day in the same conditions (“unstimulated HSPCs”)^43,42^.

Edited and stimulated (5×10^6^) and edited and unstimulated (4×10^6^) HSPCs were injected into six NSG mice (three mice with stimulated cells and three mice with unstimulated cells), of which two in each group survived to final sacrifice. Engraftment was monitored by FACS analysis of blood draws at five and eight weeks post-injection. Final engraftment was assessed at sixteen weeks post-injection, when mice were sacrificed and bone marrow (BM) and spleen cells were harvested and subjected to FACS-based lineage analysis (Fig **4A**, **S11**-**S12**). The engraftment of human cells was higher in mice injected with unstimulated HSPCs at weeks 5 and 8 than in mice injected with stimulated HSPCs, but were more comparable in BM extracted at sacrifice after 16 weeks (Fig **4A**,Fig. **S11**). Notably, substantial human CD45+ cells were detectable at 16 weeks, demonstrating the long-term repopulating potential and "stem-ness” of the original engrafted cells. Within BM, engrafted human CD45+ cells were primarily B cells (CD3+), although cells engrafted in mice injected with stimulated cells also had a substantial population of myeloid (CD33+) cells (Fig. **S12**).

**Figure 4.**
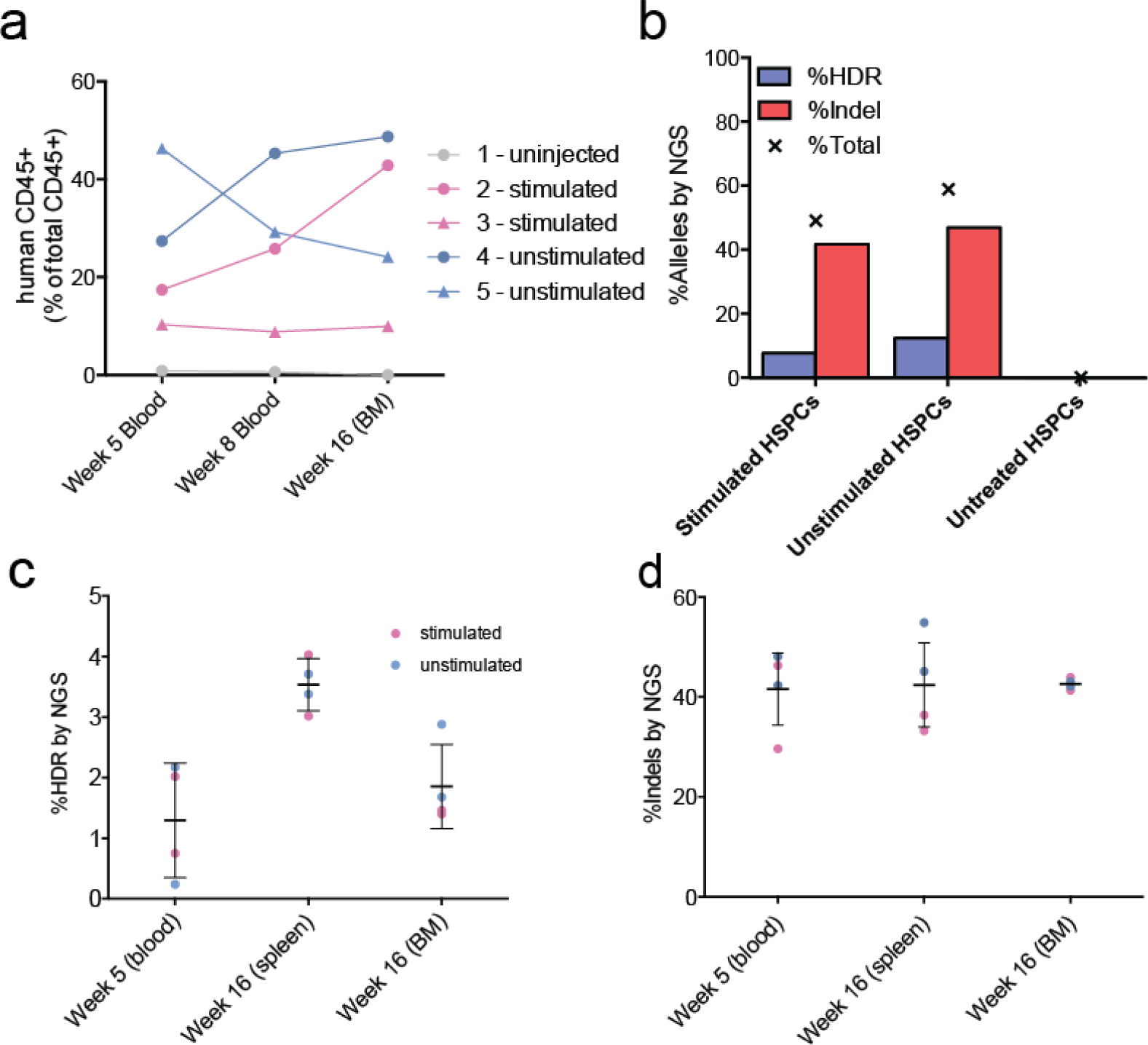
Engraftment of edited HSPCs into NSG mice. A) Plot depicting the course of engraftment of human CD45+ HSPCs cells in NSG mice injected with edited HSPCs, stimulated with expansion cytokines as indicated, compared to an uninjected mouse. B) Analysis by NGS of editing at the SCD SNP in cells prior to engraftment. C) Indel formation (left) and D) HDR-mediated editing (right) at the SCD SNP in human cells engrafted in mouse blood, spleen, and bone marrow (BM), at 5 and 16 weeks post-injection. Error bars indicate mean ± s.d. over four mice.

Genotyping of human HSPCs was assessed by NGS immediately after editing and prior to injection (Fig. **4B**), and from mice at weeks 5 (blood) and 16 (BM and spleen, Fig. **4C**-**4D**). NGS analysis revealed consistently high levels of indel alleles (42.1 ± 1.1% in bone marrow, mean ± s.d., n = 4 mice), along with maintenance of clinically relevant levels of HDR-mediated editing at the SCD SNP in both stimulated and unstimulated cells throughout the lifetime of the xenograft in both BM and spleen (1.9 ± 0.7% in bone marrow, 3.5 ± 0.4% in spleen, mean ± s.d.., n = 4 mice). These levels of long-term HDR-mediated editing are more than 9-fold greater than previously reported editing of the SCD SNP in HSCs using ZFN mRNA electroporation and ssDNA donors^18^. Since even low-level donor chimerism after allogeneic HCT for SCD can elicit a strong clinical benefit^15-17^, our results suggest that the Cas9 RNP can achieve clinically beneficial levels of SCD editing in long-term repopulating stem cells, when co-delivered with an appropriate ssDNA donor.

## Discussion

The treatment of genetic blood diseases through gene editing to replace an endogenous sequence is a long-standing goal of regenerative medicine. Unlike gene transfer by integrating viral vectors, where regulation of the introduced gene may be compromised and endogenous genes may be disrupted, gene editing corrects the disease mutation at the endogenous locus. Correction requires methods for efficient, HDR-mediated scarless editing of endogenous genes in repopulating HSCs, especially if the edit does not confer a selective advantage, such as correction of the SCD mutation^11,18^.

Here, we used the Cas9 RNP and short ssDNA donors to develop methods that efficiently edit the SCD SNP. We iterated combinations of Cas9 RNPs and ssDNA donors to edit the HBB gene in K562 cells, and then translated these reagents to human HSPCs, demonstrating efficient ***in vitro*** HDR-mediated editing of the SCD SNP with minimal off-target activity towards other genes. We further show that corrected SCD HSPCs produce substantial levels of wild-type adult and fetal hemoglobin protein, which are both protective against sickling. Importantly, HDR-mediated gene edits can be maintained ***in vivo*** in murine xenografts at levels that could be clinically beneficial (∼2% average HDR-mediated editing 16 weeks post-engraftment).

Observations of stable mixed chimerism after allogeneic HCT for SCD support the prediction that correcting the sickle allele in ≥2% of HSCs will translate into a clinically meaningful increase in levels of non-sickling circulating RBCs^15-17^. In these instances of mixed chimerism, even a small minority of donor cells in the HSC compartment is sufficient for replacement by donor erythrocytes in the blood as a consequence of improved RBC lifespan and more effective erythropoiesis. Individuals who develop stable donor-host chimerism are indistinguishable from those with full donor engraftment, including an absence of clinical events. In some patients, even 2% donor chimerism is sufficient to confer significant benefit to SCD patients^17^. Since corrected wild type alleles are dominant over SCD or indel alleles, the advantage conferred by the levels of long-term sequence correction we observe in HSCs may be more beneficial than equivalent levels of mixed chimerism ^14-16^. It is also possible that the increased levels of fetal hemoglobin protein after SCD allele editing observed here could provide further benefit, though this remains to be tested ***in vivo.***

Since endogenous sequence replacement in HSCs has historically been challenging, many attempts to cure SCD have instead relied on indirect approaches, for example by knocking out Bcl11A, a ***trans***-acting repressor of fetal hemoglobin, or by knocking out a Bcl11A erythroid-specific enhancer^26,44^. Bcl11A knockout is lethal in mice, possibly due to the repressor’s numerous cellular roles, including essential roles in neuronal and lymphoid development^45,46^. Targeting the erythroid-specific enhancer reduces these unintended effects, but this approach has yet to be fully validated. By contrast, gene correction via sequence replacement is by definition ***cis-***acting, and off-target effects are due only to the editing nuclease itself, whose lifetime in the target cell population is brief.

Although the levels of sequence correction we observe in long-term engrafted HSCs are relatively high, they are 3-5-fold lower than that measured during ***in vitro*** correction of CD34+ HSPCs. This decrease in editing frequency in repopulating HSCs, which has been observed previously^11,18^, is an impediment to the clinical utility of gene editing for diseases requiring high levels of sequence correction. It may be that HDR-edited HSCs do not engraft as well as unedited HSCs, perhaps due to toxicity of the HDR donor. Longer-term culture after editing, perhaps in the presence of recently-described HSC expansion agents, could allow edited cells to recover their ability to engraft^47,48^. Long-term re-populating HSCs may be intrinsically more difficult to edit than other progenitor cells in the CD34+ HSPC population. This deficit could be ameliorated by optimizing editing conditions using lengthy ***in vivo*** endpoints instead of the ***in vitro*** endpoints used here. This is expensive and time consuming, but may be warranted if increased editing frequencies are necessary.

In this study, we paired the Cas9 RNP with short ssDNA donors to edit longterm repopulating HSCs. This is in contrast to pairing ZFN mRNA with ssDNA donors, which has previously failed to induce substantial levels of SCD editing in HSCs^18^. Similarly, knockout in HSPCs by Cas9 mRNA delivery requires chemical protection of the sgRNA, but we found that RNP delivery yields efficient knockout even with unmodified sgRNAs^24^. It is possible that delivery of pre-formed RNP complex improves editing efficiency by remedying the multi-body problem linking Cas9, sgRNA, and HDR donor^27,28^.

Recent hematopoietic gene editing efforts have focused on viral HDR donors, often delivered at high multiplicity of infection (MOI)^12,13,18^. Advances in viral delivery technology have greatly improved safety and viral donors are a valid option for gene correction. Off-target integration of viral donor sequence remains a concern however, and high levels of off-target integration have been observed in HSCs^49,50^. Furthermore, the intensive engineering and relatively slow workflow required to design and implement viral donor delivery restricts intensive optimization of this approach and limits its eventual application to patient-specific SNPs. By contrast, a co-delivered RNP/ssDNA approach is non-viral and highly modular; development, optimization, and safety assessment are rapid. If married with anticipated developments in improving editing in re-populating HSCs, this approach may form the basis of an autologous HCT therapy for SCD and other hematopoietic diseases.

## Methods

***Synthesis of Cas9 RNPs.*** Cas9 RNP component synthesis and assembly was carried out based on published work^27^. Cas9 was prepared by the UC Berkeley Macro Lab using a published protocol^27^. Cas9 was stored and diluted in sterile-filtered Cas9 Buffer (20 mM HEPES pH 7.5, 150 mM KCl, 1 mM MgCl_2_, 10% glycerol, 1 mM TCEP). TCEP was added to storage buffer only. sgRNA was synthesized by assembly PCR and ***in vitro-***transcription. A T7 RNA polymerase substrate template was assembled by PCR from a variable 59 nt primer containing T7 promotor, variable sgRNA guide sequence, and the first 15 nt of the non-variable region of the sgRNA (T7FwdVar primers, 10 nM, Supplementary Table 1), and an 83 nt primer containing the reverse complement of the invariant region of the sgRNA (T7RevLong, 10 nM), along with amplification primers (T7FwdAmp, T7RevAmp, 200 nM each). These primers anneal in the first cycle of PCR and are amplified in subsequent cycles. Phusion highfidelity DNA polymerase was used for assembly (New England Biolabs, Inc.). Assembled template was used without purification as a substrate for ***in vitro*** transcription by T7 RNA polymerase using the HiScribe T7 High Yield RNA Synthesis kit (New England Biolabs, Inc.). Resulting transcriptions reactions were treated with DNAse I, and purified either by treatment with a 5X volume of homemade SPRI beads (comparable to Beckman-Coulter AMPure beads) or the Qiagen RNeasy purification kit, and elution in DEPC-treated water. sgRNA concentration was determined by A260 using a Nanodrop spectrophotometer (Life Technologies, Inc). Cas9 RNP was assembled immediately prior to electroporation of target cells (see below). To electroporate a 20μL cell suspension (150,000-200,000 cells) with Cas9 RNP, a 5 μL solution containing a 1.2-1.3X molar excess of sgRNA in Cas9 buffer was prepared. A 5 μL solution containing 100–200 pmol purified Cas9 in Cas9 buffer was prepared and added to the sgRNA solution slowly over ∼30 seconds, and incubated at room temperature for >5 minutes prior to mixing with target cells.

***Editing HBB in K562 cells.*** Authenticated K562 cells were obtained from the UC Berkeley Tissue Culture facility. Cells were confirmed to be free of mycoplasma contamination prior to use, using the Lonza MycoAlert kit. K562 cells were cultured in IMDM with 10% FCS, penicillin-streptomycin (100 units/μL and 100 pg/μL), plus 2 μM GlutaMax. K562 cells were edited by electroporation using the Lonza 4d nucleofector and manufacturer’s protocols (Lonza, Inc.). For each electroporation, 150,000-200,000 late log-phase K562 cells were pelleted (100 x g, 5 minutes) and re-suspended in 20μL Lonza SF solution. 20 μL cells, 10 μL Cas9 RNP containing the desired guide (see above, and Supplementary Table 1), and 1 μL 100 μM ssDNA template programming the desired edit at the SCD SNP were mixed and electroporated using the recommended protocol for K562 cells. After electroporation, K562 cells were incubated for 10 minutes in the cuvette, transferred to 1 mL of K562 media, and cultured for 48–72 h prior to genomic DNA extraction and genotyping.

***Editing HBB in primary human CD34+ HSCs.*** Cryopreserved WT human mobilized peripheral blood CD34+ HSPCs were purchased from Allcells, Inc. SCD CD34+ HSPCs were prepared by Allcells Inc. from whole blood obtained from SCD patients undergoing exchange transfusion at Benioff Children’s Hospital Oakland, and cryopreserved. To edit HSCs, ∼1 million HSPCs were thawed and cultured in StemSpan SFEM medium supplemented with StemSpan CC110 cocktail (StemCell Technologies) for 24 h prior to electroporation with Cas9 RNP. To electroporate HSPCs, 100,000-200,000 were pelleted (200 xg, 10 minutes) and resuspended in 20 μL Lonza P3 solution, and mixed with 10 μL Cas9 RNP and 1 μL 100 μM ssDNA template programming the desired edit. This mixture was electroporated using the Lonza 4d nucleofector and either of two protocols (“1”: DO100, "2”: ER100). Electroporated cells were recovered in the cuvette with 200 μL StemSpan SFEM/CC110 for 10–15 minutes, and transferred to culture in 1 mL StemSpan SFEM/CC110 for 48 hours post-electroporation. Half of the cells were removed for genotyping ("un-expanded HSPCs”) while the remaining cells were transferred to erythroid expansion media (StemSpan SFEM II with StemSpan Erythroid Expansion supplement, [StemCell Technologies]) for 5 additional days prior to genotyping of expanded cells.

***Editing HSPCs prior to injection in NSG mice.*** For each mouse, 750,000-1,000,000 CD34+ HSPCs were edited. Editing was performed using a scaled-up reaction volume from ***in vitro*** experiments above. 750,000-1,000,000 HSPCs were thawed and recovered for 24 h prior to editing. Both stimulated and unstimulated conditions used StemSpan SFEM medium supplemented with StemSpan CC110 cocktail, stimulation was for 3 days while unstimulated was as above. Prior to editing, HSPCs were pelleted and resuspended in 100 μL P3. 500 pmol Cas9 RNP was prepared in 50 μL Cas9 buffer. RNP, cells, and 5 μL 100 μM donor template were mixed in a large-sized cuvette and electroporated using the Lonza 4d Nucleofector and protocol ***“2"*** (ER100). Cells were recovered by addition of 400 μL StemSpan SFEM/CC110 to the cuvette, prior to culture in recovery media for 24 hours at a density <1 million cells/mL.

***Xenografting of human CD34+ HSPCs into NSG mice.*** NSG mice (JAX) were maintained in clean conditions. 7 week old female mice were subjected to 2.5Gy x-irradiation 4 hours prior to tail vein injection of edited cells under isoflurane anesthesia. At 5 weeks and 8 weeks after injection, 200pl blood was obtained from the submandibular vein under isoflurane anesthesia. 16 weeks after injection, mice were euthanized, and bone marrow and spleen were recovered for analysis.

***Flow cytometry.*** Cells were prepared from peripheral blood, bone marrow, or spleen of NSG mice by standard methods, stained with antibodies to the indicated cell surface markers, and analyzed on a BD FACS Fortessa flow cytometer. Flow cytometry data was analyzed using the Flowjo software package.

***Differentiation of HSCs into erythroblasts.*** After electroporation, cells were recovered and placed in StemSpan SFEM/CC110 for 24 hours. They were then transferred to StemSpan SFEM II with StemSpan Erythroid Expansion supplement and grown for 7 days with maintenance of optimal density (200,000-1,000,000 cells/mL). The resulting erythroid progenitors were transferred to StemSpan SFEM II with 3U/ml erythropoietin (Life Technologies), 3% normal human AB serum (Sigma), and ΙμΜ mifepristone (Sigma). They were then cultured for a further 5 days with daily monitoring of cell morphology after Wright-Giemsa staining; at the conclusion the majority of the cells were enucleated. Cells were then lysed in hemolysate Reagent (Helena Laboratories) for preparation of hemoglobin for HPLC.

***Genotyping of edited cells.*** Pools of edited cells (K562 cells, CD34+ HSCs, and enucleated cells from mouse blood) were lysed and their genomic DNA extracted using QuickExtract solution (Epicentre Inc.) to a final concentration of ∼5,000 haploid genomes/μL. A 350 bp region around the SCD SNP and Cas9 cut site was amplified by PCR using Q5 DNA polymerase (New England Biolabs, Inc), and primers 1F and 1R (see Supplementary Table 1). NHEJ-mediated indel formation within each pool was estimated by T7 endonuclease I digestion using manufacturer’s protocols (New England Biolabs, Inc). HDR-mediated editing was assessed by restriction digest with either ***SfcI*** (for WT->SCD edits) or ***Hpy 188111*** (for SCD->WT edits). For analysis of editing by next-generation sequencing (NGS), a second, low-cycle PCR using primers 4F and 1R was used to generate amplicons of an appropriate length, prior to Illumina TruSeq adaptor ligation and purification (Bioo Scientific DNAseq kit or Illumina TruSeq Nano HT kit). To avoid contamination of ssDNA donor sequence, 1F and 1R amplify outside the genomic region matching donor, and 4F does not anneal to the ssDNA donor. Libraries from 12–96 pools of edited cells were pooled and run on a single MiSeq lane, using a paired-end 150 cycle read. HDR-mediated editing of the SCD SNP was also assessed by droplet digital PCR (ddPCR, QX200, Bio-Rad Laboratories, Inc.). ddPCR assays (1x assay: 900 nM primers, and 250 nM each probe) used TaqMan probes specific for unedited and HDR-edited alleles, with one primer positioned outside of the template matching region of HBB to prevent amplification of donor template. Assays were run using ddPCR supermix for probes (no dUTP) with the following thermal cycling protocol: 1)95°C 10 min; 2), 94°C 30 s; 3), 55°C 1 min; 4), 72°C 2 min, 5) repeat steps 2–4 39 times; 5), 98°C 10 min with all the steps ramped by 2°C/s.

***NGS Data analysis.*** 20 million MiSeq reads were de-multiplexed and analyzed using a custom analysis workflow written in Python. Each sample contained >75,000 reads, generally much more. For each sample, reads were called as “indel”, “HDR”, or “unedited”. Any read containing an indel within a window of 12–16 bases around the predicted cut site was called as “indel”, and remaining reads were called as “unedited” or “HDR” based whether they matched either the unedited sequence or the ssDNA donor sequence at the SCD SNP.

***HPLC analysis of edited SCD HSCs.*** HPLC analysis was performed as previously described^18^. Briefly, edited SCD HSPCs differentiated into erythroblasts were harvested and lysed in Hemolysate reagent (Helena Laboratories). Cell lysates were characterized by HPLC (Infinity 1260, Agilent) using a weak cation-exchange column (PolyCAT A^™^, PolyLC_INC_.)⋅⋅ Analysis and peak integration was performed using OpenLAB CDS Chemstation software. FASC Reference Material (Trinity Biotech) was used to define the elution time of common hemoglobins (HbF, HbA, HbS, and HbA2).

**Figure S1.**
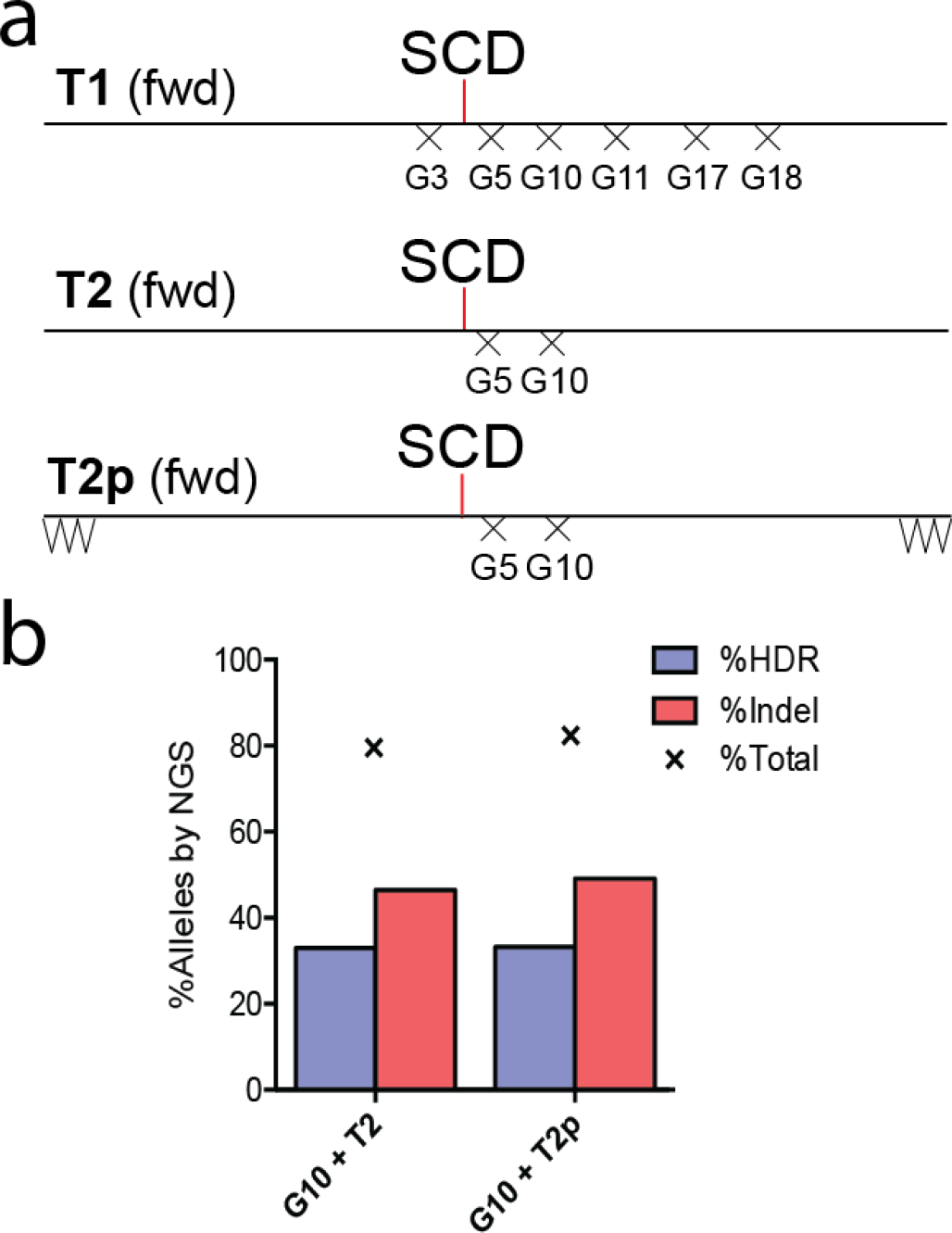
ssDNA donor templates used with various sgRNAs for editing of K562 cells with the Cas9 RNP, effect of ssDNA template protection on editing K562 cells. A) Initial templates are 195 nt long, and edit the WT SNP to SCD. To prevent re-cutting, template 1 encodes silent mutations for the PAMs of all the sgRNAs tested. Templates **T2** and **T2p** have PAM muations only for **G10** and **G5.** Template **T2p** has three 3’ and three 5’ phosphorothioate linkages to prevent degradation by endogenous exonucleases. B) Editing outcomes of K562 cells edited with unprotected template **T2** and phosphorothioate-protected **T2p**, analyzed by NGS.

**Figure S2.**
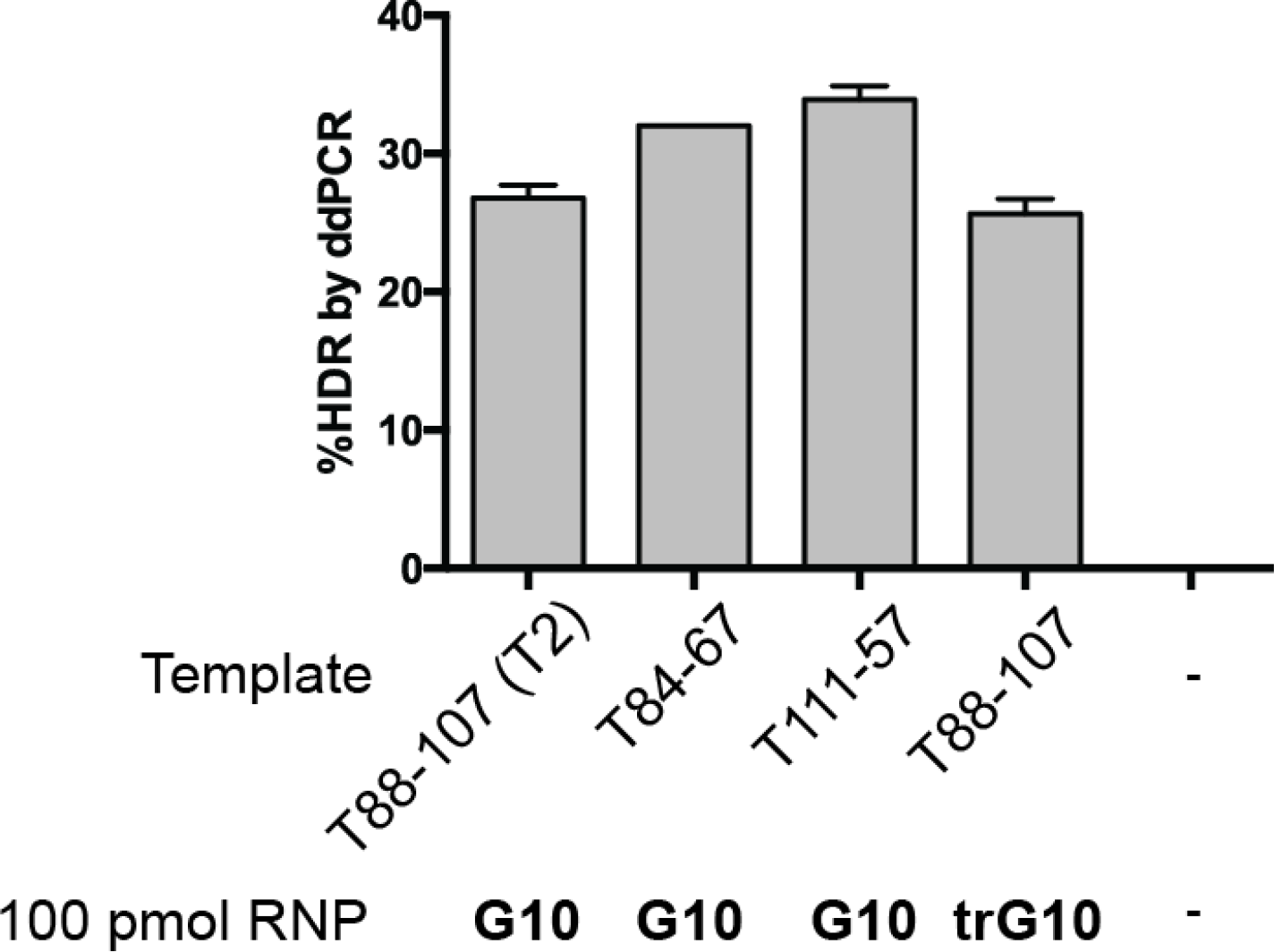
Verification of HDR-mediated editing in K562 cells by droplet digital PCR (ddPCR). Genomic DNA from K562 cells edited with indicated RNPs and templates were analyzed by ddPCR using probes and primers as indicated in Materials and Methods. Error bars are SEM of 3 independent biological replicates, each consisting of 3 technical replicates. Template **T88-107** is identical to **T2** (Fig. S1)

**Figure S3.**
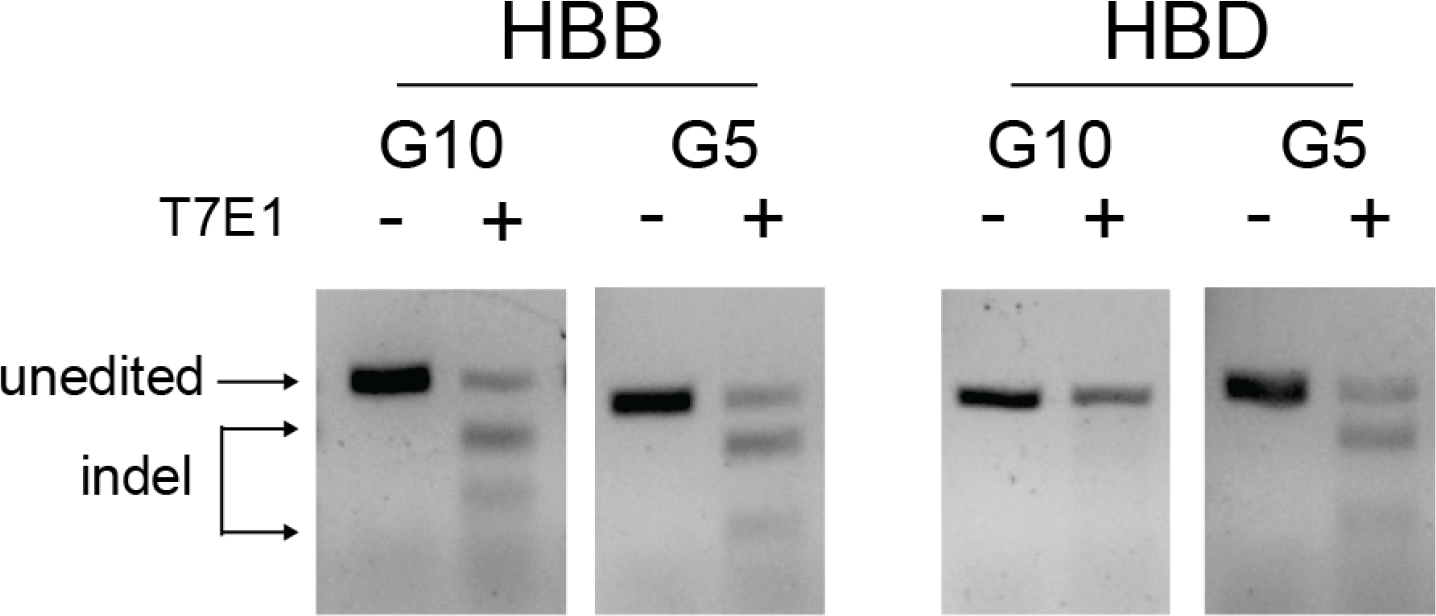
The G5 RNP induces indels at HBD. **T7 endonuclease 1 assay** of PCR amplicons from the HBB SCD region or the corresponding region in HBD, from K562 cells edited with indicated RNPs, analyzed on a 2% agarose gel. Both the **G5** and **G10** RNPs cut efficiently at HBB, and **G5** also cuts at HBD.

**Figure S4.**
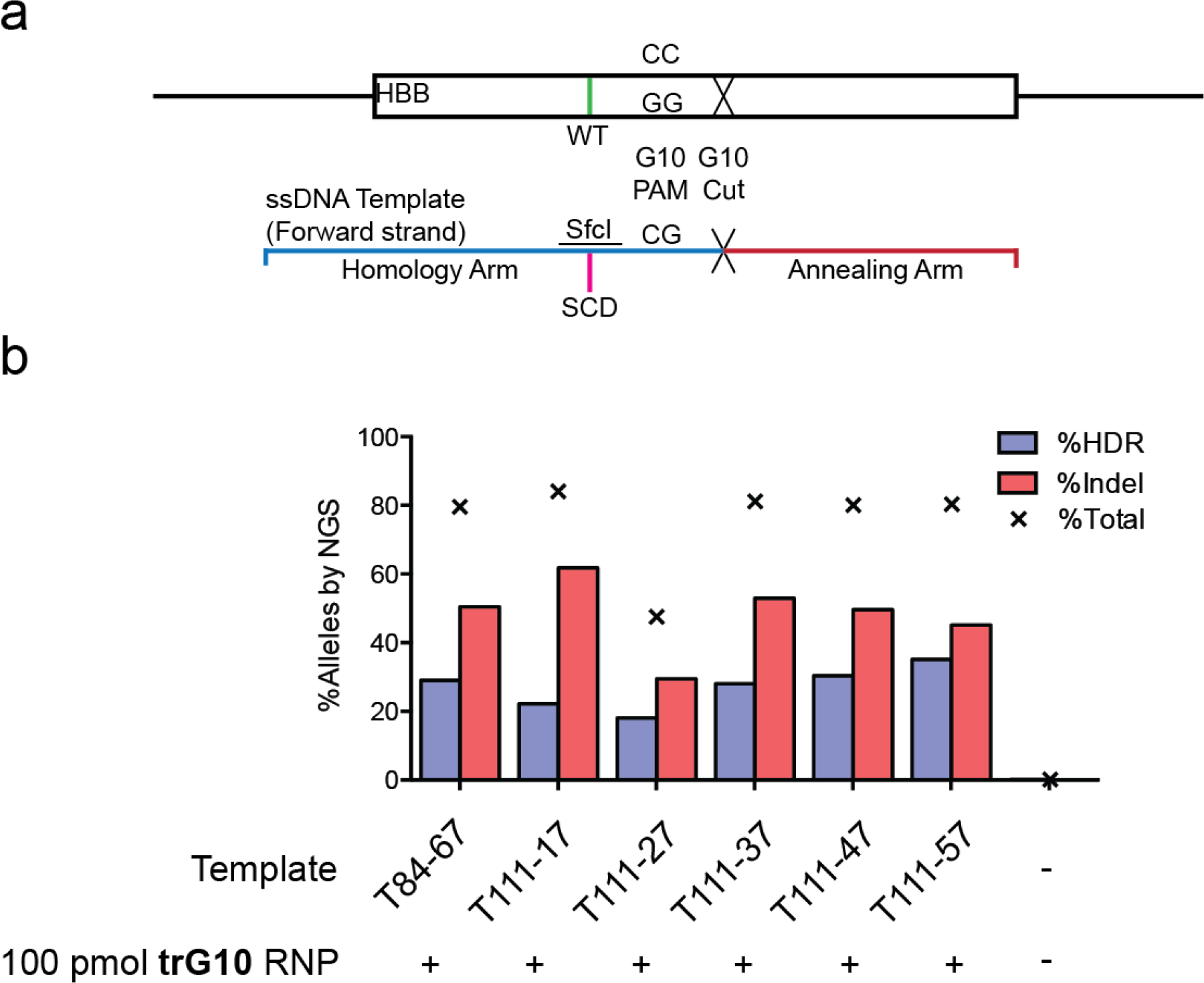
Template optimization in K562 cells, analyzed by NGS. A) Design of ssDNA templates, relative to the G10 cut site. Templates consist of a shorter “annealing” arm that anneals to the strand liberated by RNP binding, and a longer “homology” arm that drives incorporation of the desired edit. B) K562 cells were electroporated with the **trG10** sgRNA as indicated, along with the indicated asymmetric template, defined by the lengths of the homology and annealing arms, respectively (homology length-annealing length). Editing outcomes were assessed byNGS.

**Figure S5.**
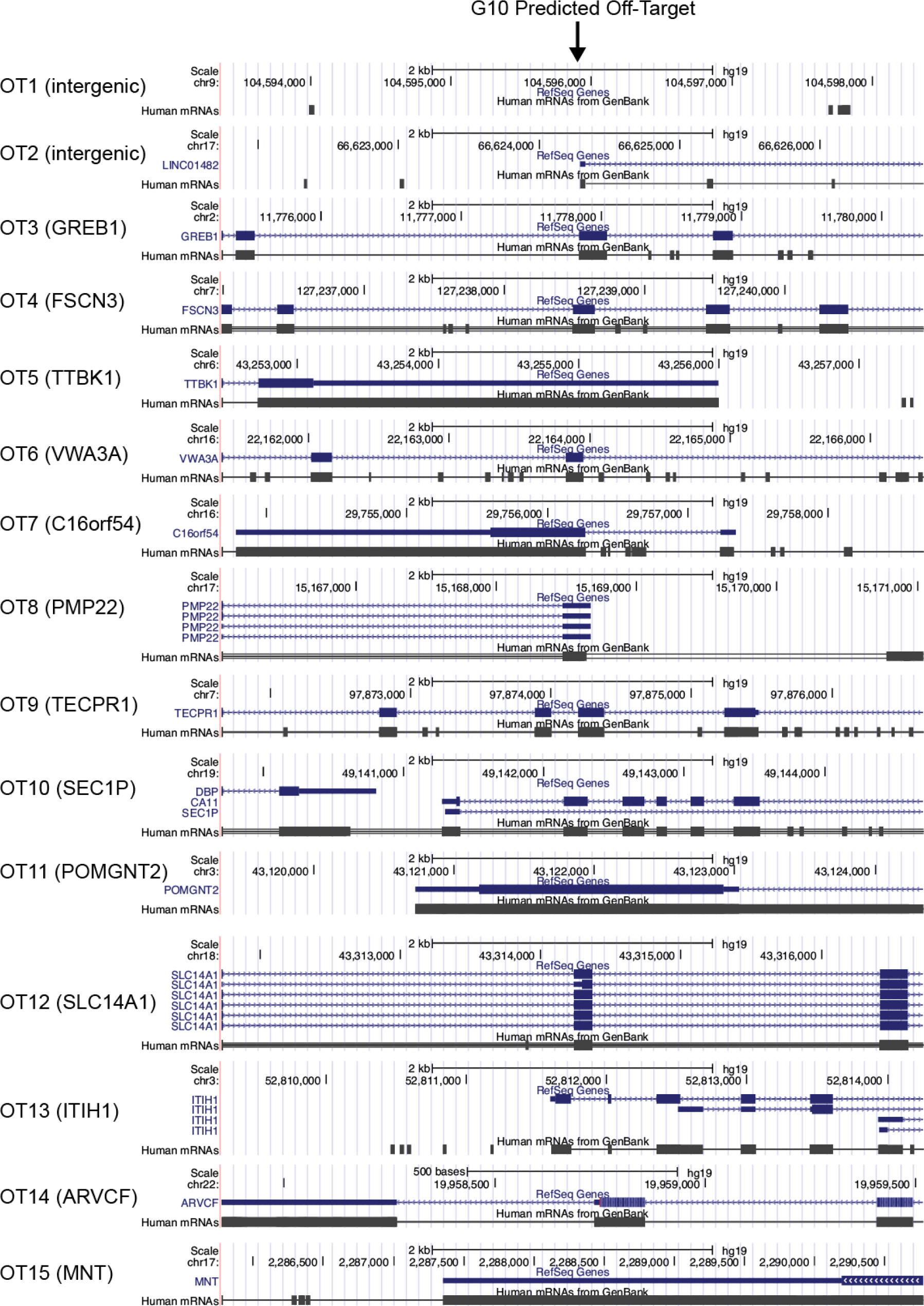
Genomic context of predicted off-target cut sites for the G10 RNP. Sites were selected using the online CRISPR-design tool, according to criteria discussed in the text. Screenshots of the 5 kb region near the off-target site were prepared using the UCSC genome browser.

**Figure S6.**
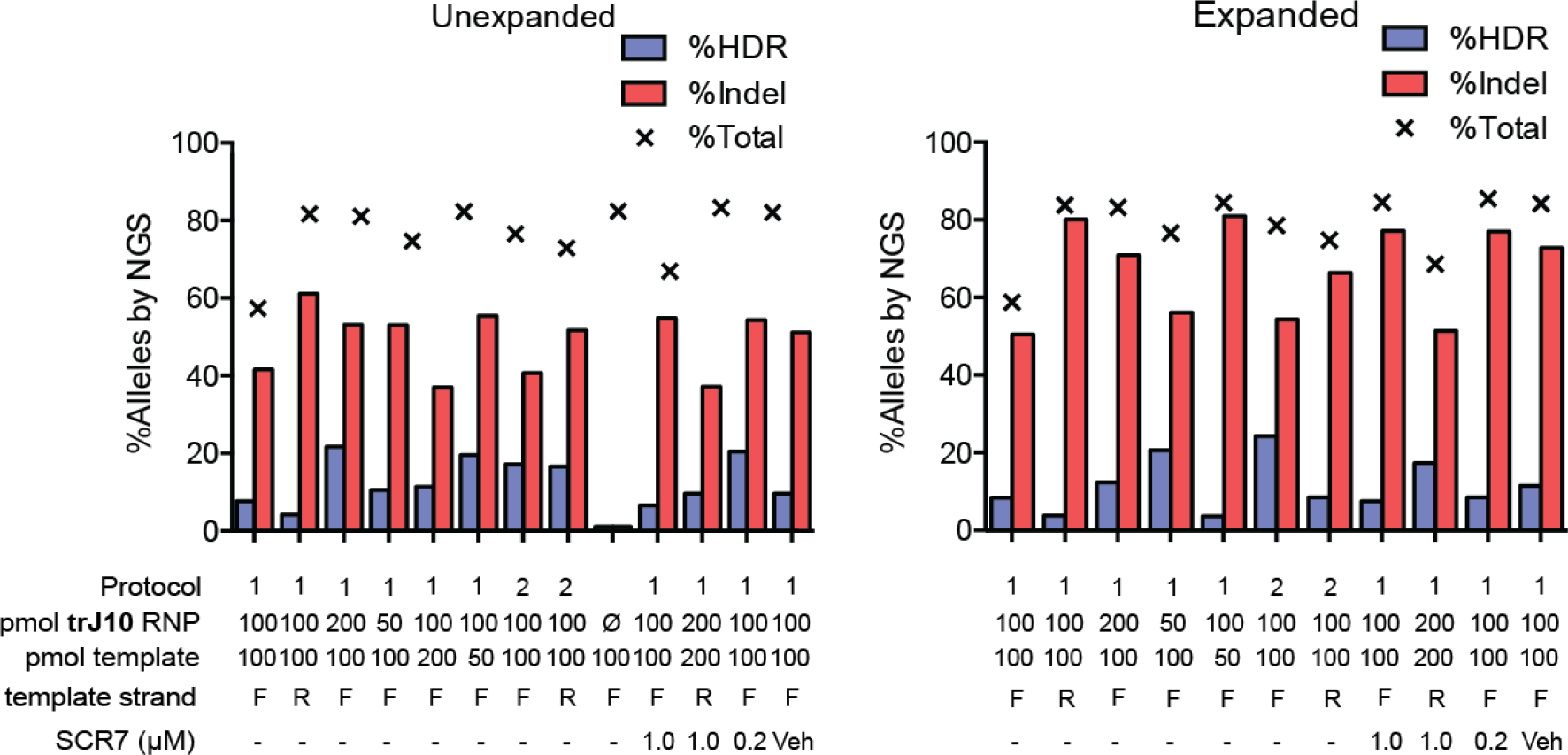
Optimization of editing conditions in HSPCs using the trJlO RNP. Editing outcomes of un-expanded HSPCs (cultured for 2 days after editing in HSC expansion medium) and erythroid-expanded (cultured for 5 additional days in erythroid expansion medium) HSPCs after electroporation with the indicated doses of templates, **trJG0** RNP, and Scr7 (NHEJ inhibitor), and either protocol 1 (Lonza DOIOO) or protocol 2 (Lonza ER100). Either forward-strand matching (F) or reverse strand-matching (R) templates (template T2, and its reverse complement, Fig. S1A, Supplementary Table 1) were provided as indicated.

**Figure S7.**
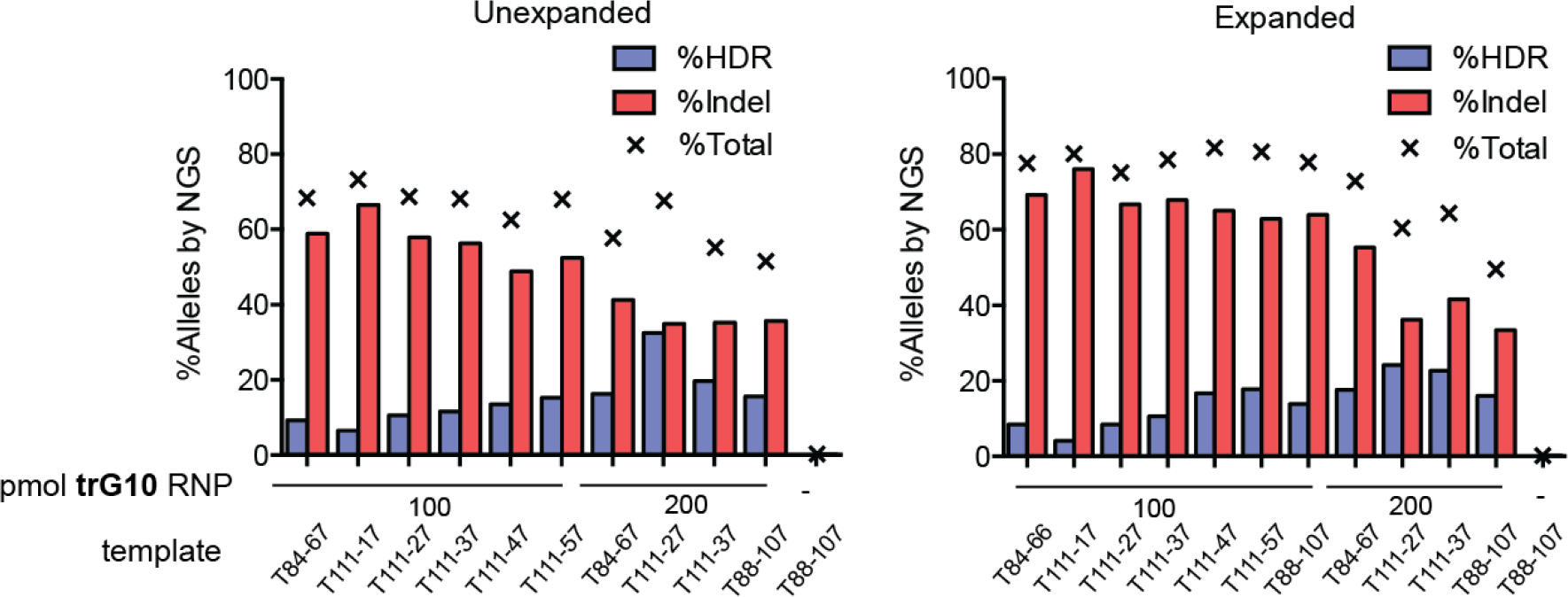
Template optimization in HSPCs. Editing outcomes of HSPCs cultured non-expanding and erythroid-expanded conditions, assessed by NGS. Templates are designed and described as for K562 cells (Fig. S4). 150,000 CD34+ HSPCs were edited with indicated templates and RNPs as indicated, and protocol 2 (Lonza 4d code ER100.

**Figure S8.**
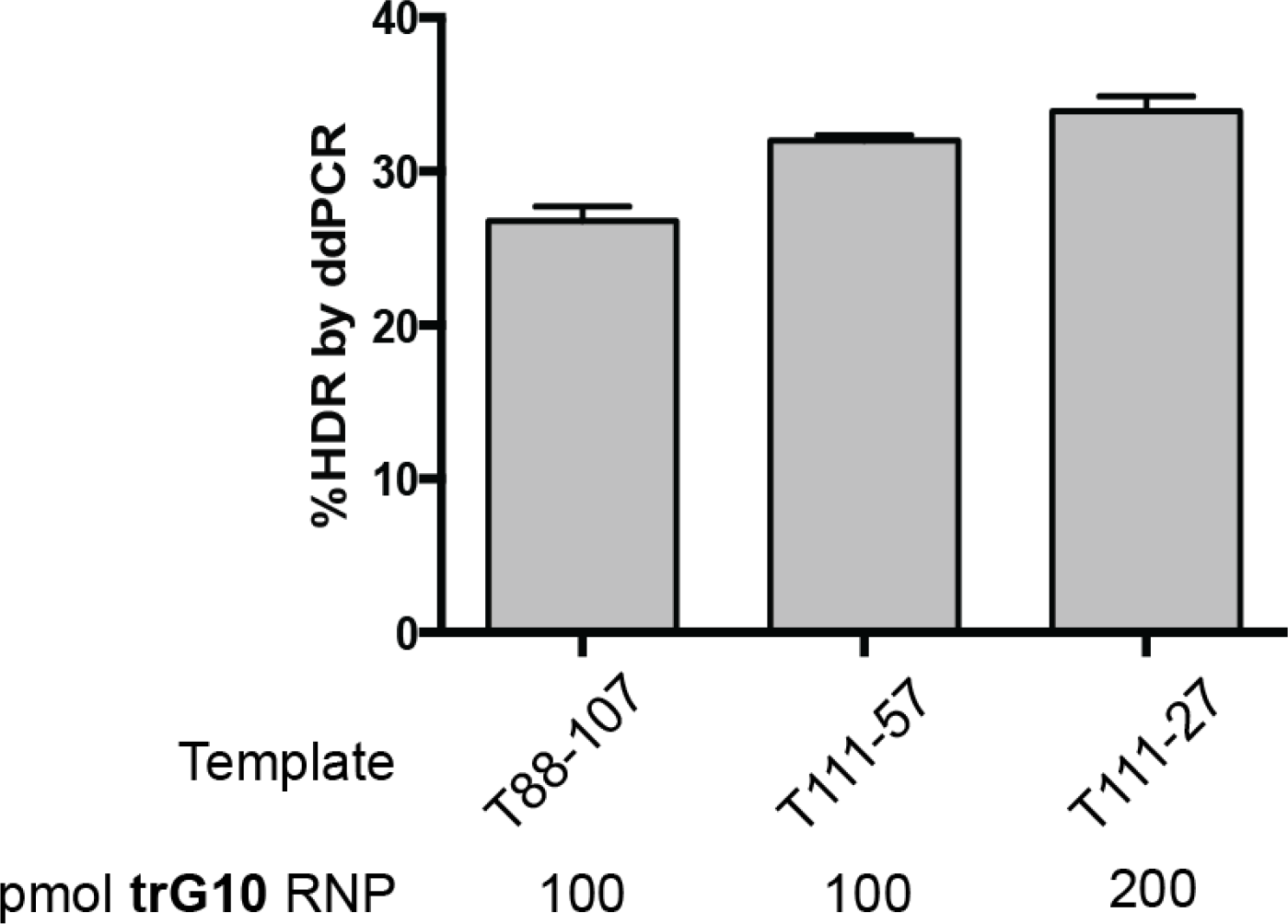
Editing in WT un-expanded HSCs in biological replicate by ddPCR. HSCs (n=3 biological replicates) were edited with the indicated doses of **trG10** RNP, along with the indicated asymmetric template, and cultured in non-expanding conditions for 2 days prior to genotyping by ddPCR. HDR-mediated editing was quantified by ddPCR of genomic DNA extracted from edited cells, using probes for WT (unedited) and SCD (edited) alleles. Error bars indicate the SEM of the means from three technical replicates of three biological replicates, from a single healthy donor.

**Figure S9.**
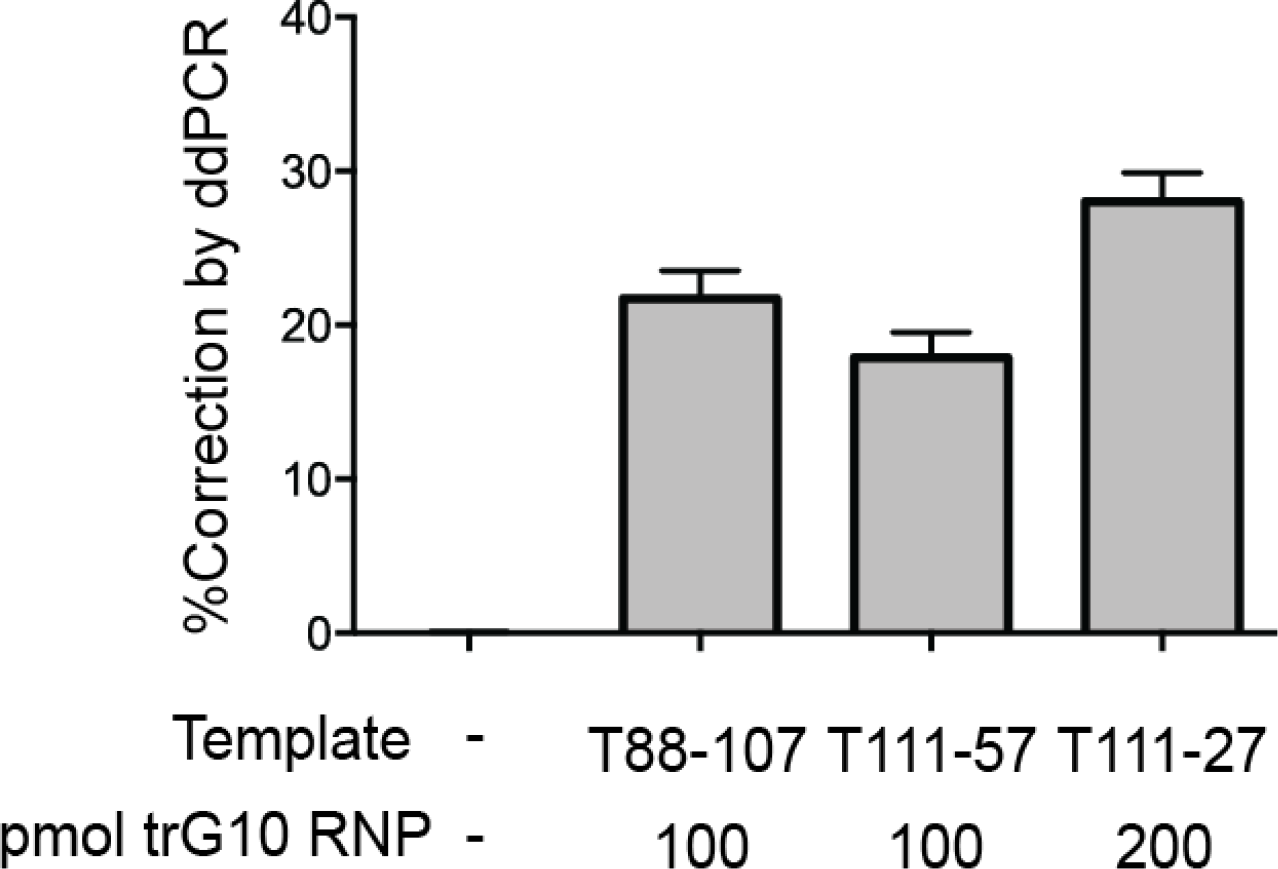
HDR-mediated correction of SCD HSPCs, assessed by ddPCR. HSPCs from SCD patients were edited with indicated templates and the **trG10** RNP, and cultured for 2 days in non-expanding condtions (identical to conditions in Fig. 3). Editing was then assessed by ddPCR of genomic DNA extracted from edited cells, using TaqMan probes for the SCD (unedited) and WT (edited) alleles.

**Figure S10.**
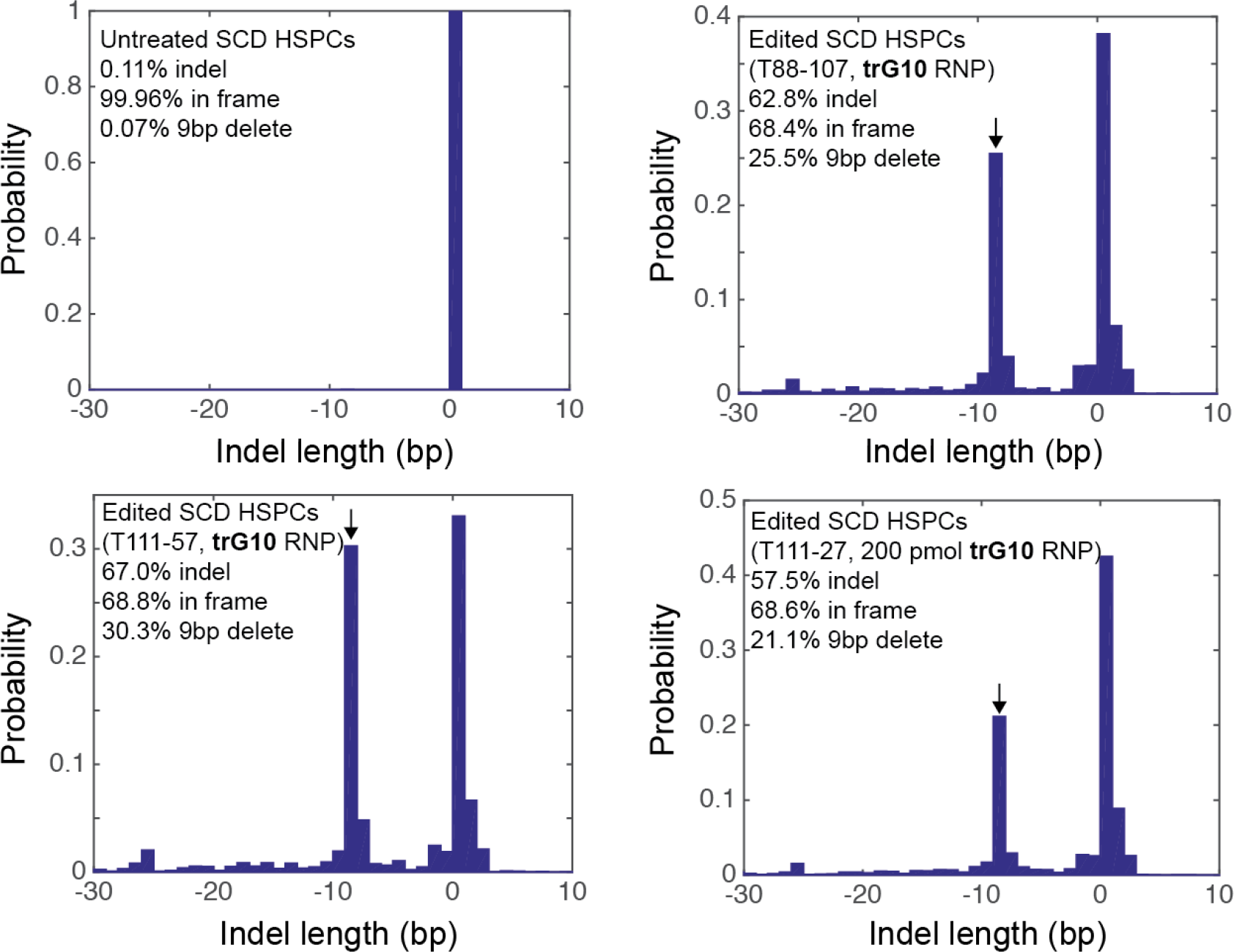
Characteristics of indels formed by the **trG10** RNP in SCD HSPCs. Histograms of indel lengths generated in cells edited with the **trG10** and the indicated asymmetric templates, compared to untreated SCD HSPCs identical to Figure 3A. WT and edited (non-indel) alleles are scored as 0 bp indels, deletions are <0 bp indels, and insertions are >0 bp indels. An in-frame 9 bp deletion is prominent in all pools of edited cells. This indel results from use of a 5 bp microhomology flanking the **trG10** cut site and is also seen with the **G10** sgRNP.

**Figure S11.**
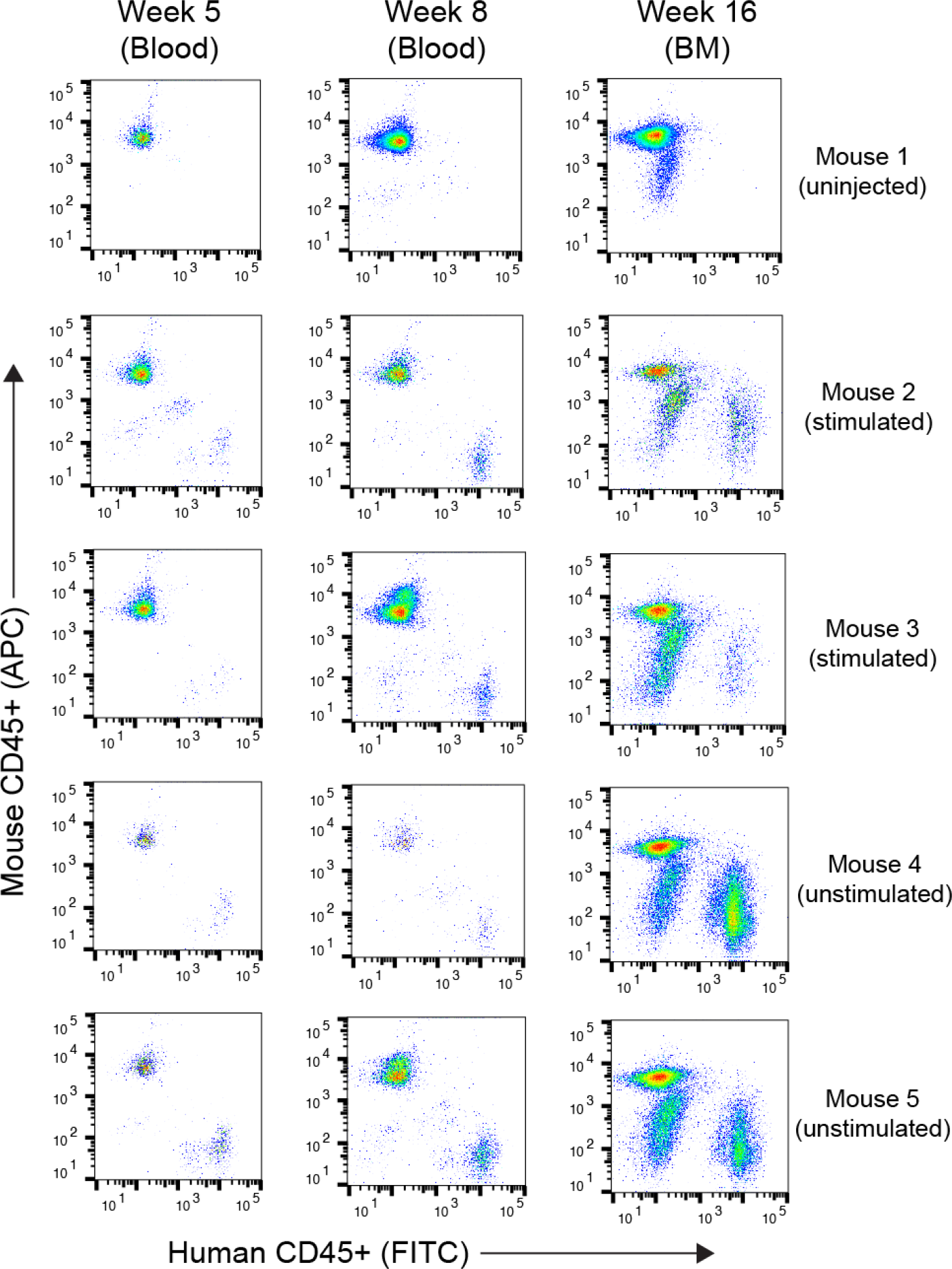
FACS plots depicting engraftment of edited HSPCs in NSG mouse. Mice were engrafted with either no cells (uninjected), edited and stimulated human HSPCs (stimulated), or edited and unstimulated human HSCs. At indicated times, engraftment in either blood or bone marrow (BM) was determined by immunostaining and FACS analysis with human anti-CD45-FITC and mouse anti-CD45-APC.

**Figure S12.**
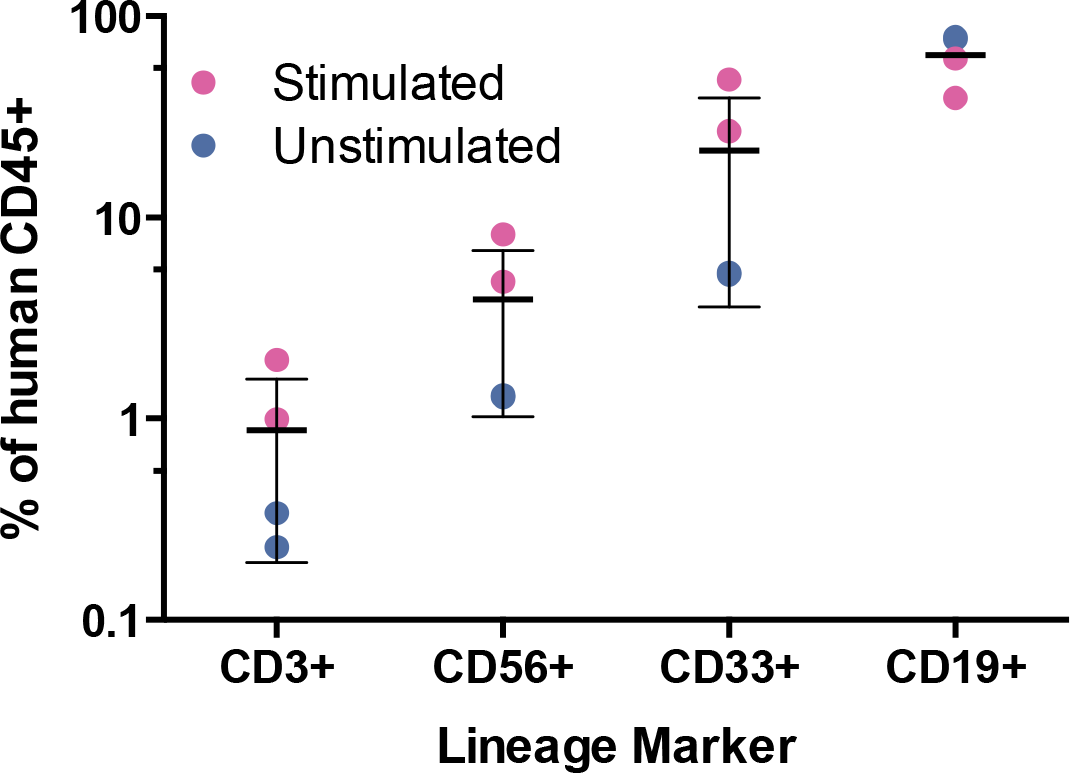
Final lineage analysis of edited HSCs engrafted in NSG mouse bone marrow, by FACS. Stimulated and unstimulated CD34+ HSPCs were edited with the **trG10** RNP, and injected into 4 mice. Bone marrow from mice 16 weeks postinjection was harvested, stained with antibodies against indicated human-specific markers along with human CD45. Error bars represent Mean ± s.d. over four mice.

